# Mapping pathogenic patterns in membrane transporters from the GLUT transporter family

**DOI:** 10.64898/2026.06.28.735151

**Authors:** Nina Kadášová, Dominik Martinát, Anna Špačková, Ivana Hutařová Vařeková, Karel Berka

## Abstract

Non-synonymous amino acid substitutions (missense mutations) are common in the general population; some are causative of serious disease. Depending on their structural context, they can disrupt protein function, folding, or dynamics. Computational predictive methods developed in recent years, such as AlphaMissense, provide new insights into how missense mutations affect protein structure by predicting and mapping their pathogenicity across each amino acid in the human proteome. In this study, we identify recurring patterns of pathogenicity prediction across the GLUT family membrane transporters encoded by genes *SLC2A1-14*. Within the GLUT transporter family, we observe higher pathogenicity profiles in the transmembrane domains, particularly in pore-lining and binding-site residues. Predicted missense pathogenicity is elevated throughout residues assigned to the central cavity, suggesting sensitivity of the transport pathway. Another finding shows higher pathogenicity in specific transmembrane helices of the protein, with the same pattern across all proteins. On the other hand, we observed lower pathogenicity values in some representatives of the GLUT family. We validated these predictions against clinically observed missense variants from ClinVar and benchmarked three prediction methods. AlphaMissense showed the strongest concordance with clinical classifications (AUC = 0.88), and the structural distribution of clinically reported pathogenic variants broadly recapitulated the predicted pathogenicity landscape. On the other hand, we observed lower pathogenicity values in some representatives of the GLUT family, and in select cases, clinical and predicted data diverged, suggesting more localized functional constraint than genome-wide prediction alone would indicate. These findings show that the pathogenicity of glucose transport within the GLUT family may be shaped by functional redundancy and physiological essentiality across GLUT groups, supported by convergent evidence from structural, computational, and clinical variant data.

## 1. Introduction

The primary structure of the protein is a linear sequence of amino acids, which provides an architectural plan for subsequent conformations, ranging from local folding (secondary structure) to complexes (tertiary and quaternary). (Stollar and Smith, 2020) The intact structural architecture is essential for proper protein function and for maintaining the organism’s homeostasis. (Cheng et al., 2023) Any changes in the primary sequence can lead to defective protein folding, reduced stability, and, finally, loss of function. Missense mutations (MMuts) are those types of disturbances in the primary sequence, where one amino acid is substituted for another. (Cheng et al., 2023; Stefl et al., 2013; Tian et al., 2023) These molecular changes are directly connected with the etiology of a wide spectrum of human diseases. (Badonyi and Marsh 2024; Stefl et al. 2013)

Our focus is on the facilitative glucose transporter family (GLUT), encoded by genes *SLC2A1-14*, comprising 14 members divided into three classes based on transported substrate, protein sequence, and structural similarity. (Holman, 2020) These proteins are structurally characteristic representatives of the major facilitator superfamily (MFS). (Long and Cheeseman 2015) All GLUT proteins contain twelve transmembrane (TM) helices arranged into two six-helix domains—the N-domain (TM1–6) and the C-domain (TM7–12)—which are pseudo-symmetrical and belong to the MFS (Figure 1). Between these two halves lies a central cavity that accommodates the substrate-binding site. (Drew et al., 2021)

**Figure 1:**
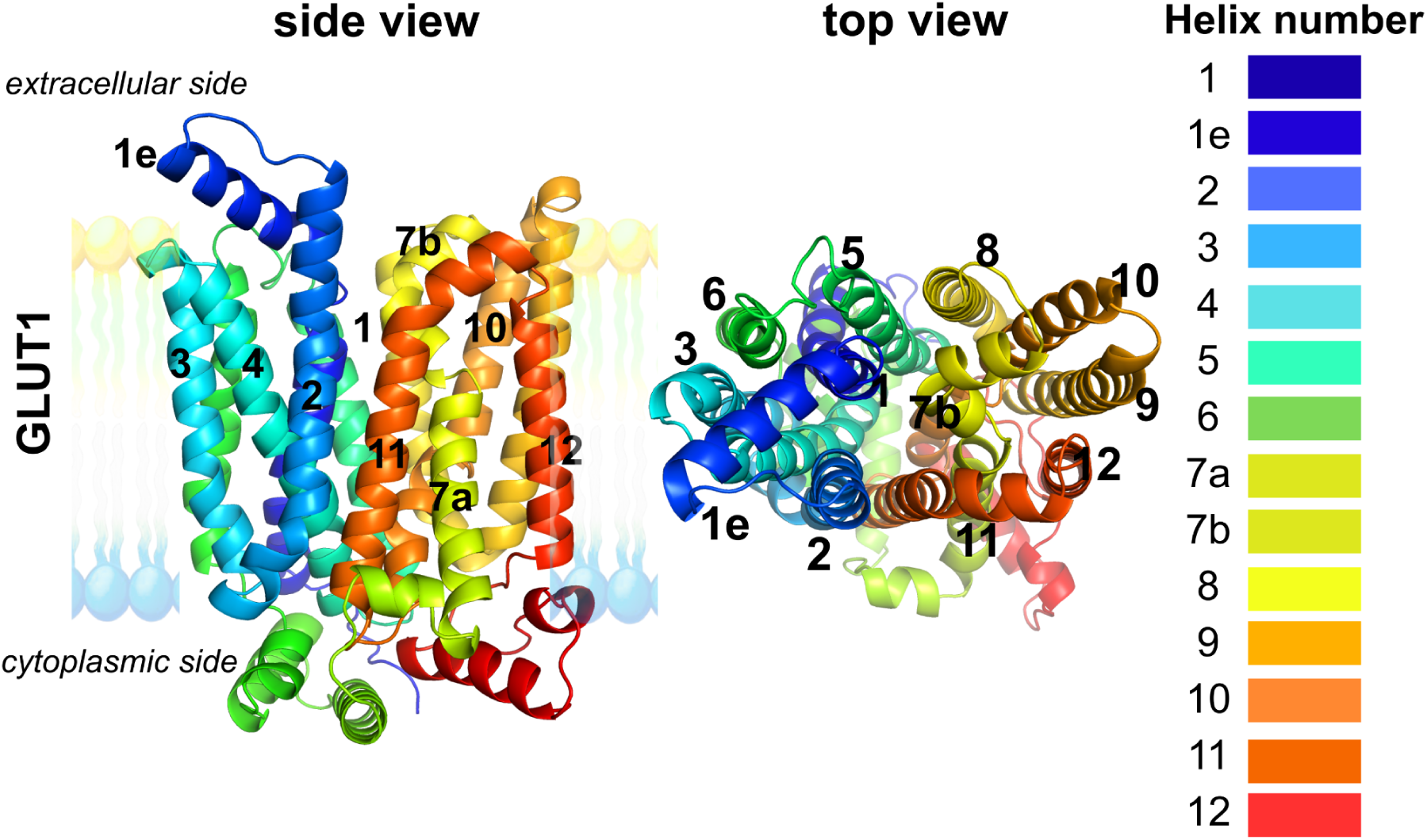
The representation of the helices in GLUT transporters is shown on GLUT1. The helices were numbered in order of their sequence within the amino acid sequence. The color scheme from blue to red follows the arrangement from the C- to the N-terminus. The protein model is embedded in membranes marked by a lipid bilayer separating the extracellular and cytoplasmic sides.

GLUTs are also key players in metabolic regulation by facilitating hexose transport across membranes. (Holman 2020; Long and Cheeseman 2015) Mutations in *SLC2A* genes, such as *SLC2A10*, are associated with clinically important states. Besides type 2 Diabetes, they are also associated with various types of cancer (Li et al., 2025; Miller et al., 2024; Ostrzygało et al., 2025; Ren et al., 2025; Zhang et al., 2022), glucose transporter-1 deficiency syndrome (Giugno et al., 2024; Mastrangelo et al., 2024), and arterial tortuosity syndrome. (Esmel-Vilomara et al., 2023) Understanding of the influence of MMuts in these structurally similar membrane proteins is paramount.

Despite substantial advances in *in silico* tools for predicting the pathogenicity of amino acid substitutions, accurately interpreting the clinical significance of missense variants remains challenging. Traditional tools like SIFT and PolyPhen-2, which primarily rely on evolutionary conservation (Flanagan et al., 2010; Garcia et al., 2022), often struggle to discriminate benign from pathogenic variants in structurally complex and conformationally dynamic regions typical of these proteins. Recent deep learning–based models, including AlphaMissense, have significantly enhanced predictive accuracy. (Cheng et al., 2023; Krenn et al., 2025; Tordai et al., 2024) However, gaps persist in linking these predictions to precise structural contexts relevant to function. Applying predictive techniques such as Alphamissense, PolyPhen-2, or SIFT to protein structure models can provide insight into sites with elevated predicted deleteriousness scores. This reveals the importance of these sites for protein function and integrity.

Previous studies indicate that areas with higher computationally predicted intolerance to substitution are not random but exhibit structural dependence. Recent data pointed out that sites with elevated predicted deleteriousness scores occur in critical functional regions of proteins, such as membrane-bound regions, interaction binding sites (IBS), mitochondrial proteins, and housekeeping genes. (Tordai et al., 2024) Another study done on the Cytochrome P450 family goes deeper into the protein structure itself. The analysis focused on tunnels identified via MOLE (Raček et al., 2025) in cytochromes P450 and found an increase in pathogenic MMuts along protein tunnels towards the cofactor-binding site. (Špačková et al., 2025) Together, these findings demonstrate the importance of protein regions within its secondary structure for its function.

This study provides a more detailed analysis of the distribution of MMuts pathogenicity hotspots within the GLUT protein structure. We aim to compare pathogenicity in the TM region, in the protein channel, and at the binding site. We also intend to compare the prevalence of pathogenic MMuts in the extracellular and intracellular domains with that in the entire protein and its TM region. Our objective is to determine whether the entire TM region exhibits the same level of pathogenicity as the amino acid residues lining the protein tunnel, which serves as the main transport pathway, or the binding site itself.

## 2. Materials and Methods

### 2.1. Protein structures

For the study, 14 protein amino acid sequences were selected from the UniProt database (The UniProt Consortium 2025) to ensure they corresponded to those of *Homo Sapiens*. Each sequence represents one of the 14 proteins in the GLUT transporter family. Nowadays, 22 structures of 5 proteins from the GLUT family (Khandelwal et al. 2025; Lee et al. 2024; Deng et al. 2014; Wang et al. 2022; Yuan et al. 2022; Matsushita et al. 2025; Zhao et al. 2026; Custódio et al. 2021; Kapoor et al. 2016) are available in the Protein Data Bank. As it is still not possible to find all GLUT representatives in the PDB structures, *in silico* models generated with the Boltz-2 tool (Passaro et al. 2025) were employed to analyze the entire GLUT family.

### 2.2. MOLEonline

The interactive, web-based application MOLE was used to analyze the pores in GLUTs. MOLE (Raček et al. 2025) is used to compute tunnels in biomacromolecular structures. In this study, its other computational capability, designed to detect transmembrane pores, was used. For each protein, we identified one computed pore as its transport pathway. In the final analysis, lining residues around the central cavity with openings, as counted by MOLE, were used. The extendable properties were set to the membrane region only, with a Probe Radius of 13 Å and an Interior Threshold of 0.7 Å.

### 2.3. PrankWeb

A search for protein binding sites was performed using PrankWeb, a web server that predicts protein–ligand binding sites from a given three-dimensional structure. (Jakubec et al. 2022; Polák et al. 2025) The PrankWeb interface uses the P2Rank standalone method (Krivák and Hoksza 2018), which leverages machine learning to identify binding sites on the protein surface. Using PrankWeb, we identified multiple binding sites, but for the study, we used only the one with the highest score and probability. This binding site was also most often predicted at the pore entrance or, in its vicinity, which suggested that it would be the main binding site near the central cavity.

### 2.4. DeepTMHMM

For transmembrane topology prediction, DeepTHMMM was used, which utilizes a deep learning encoder-decoder sequence-to-sequence model. (Hallgren et al. 2022) Output from this method is labeled as the amino acid sequence of the protein according to the corresponding protein topology. For GLUT proteins, we needed to separate the transmembrane region, which DeepTMHMM recognized as alpha-membrane (M), from the intracellular (I) /outside (O) sides of the cell.

### 2.5. DALI

DALI is a web server for comparing 3D protein structures using distance-matrix alignment. DALI supports two structure comparison modes, which are “Pairwise” and “All to all”. (Holm 2020) For this study, the second type of structural comparison was used, in which all 14 GLUT protein structures obtained by Boltz-2 were uploaded. The outcome was a structural similarity dendrogram used for structural alignment visualization.

### 2.6. OverProt

OverProt is a web-based tool for generating consensus secondary structures for protein families, displaying characteristic helices and β-sheets across multiple structures. It thus provides an overview of the family’s common architecture and its variability. (Midlik et al. 2022) This study used OverProt to annotate transmembrane helices in GLUT protein structures, thereby enabling consistent identification of helical segments within the membrane-spanning region.

### 2.6. AlphaMissense

AlphaMissense is a novel deep-learning-based prediction tool that leverages protein structure information from AlphaFold 2. (Jumper et al. 2021) For the prediction, AlphaMissense uses protein sequence and predicted structural context. Deep learning is trained on the frequency of MMuts in the population. (Cheng et al. 2023) In various studies, AlphaMissense shows better prediction abilities than previously used tools. (Krenn et al. 2025; Tordai et al. 2024; Kurtovic-Kozaric et al. 2024)

### 2.7. PolyPhen-2

PolyPhen-2 (Polymorphism Phenotyping) is an automated tool for predicting the possible effect of non-synonymous amino acids substitution on the function or structure of human proteins. It uses a machine learning classifier-based approach. PolyPhen-2 is used to predict the impact of non-synonymous amino acid substitutions on proteins by analyzing multiple sequence alignments and 3D structural features. It assumes that amino acid variations at conserved positions are more likely to cause functional changes. Amino acid variations affect stability, binding sites, solubility, and other critical parts of protein structure. For these reasons, amino acid variations can influence the entire protein’s structure. Changes caused by amino acid variations are likely to be linked to the protein’s function. This is assumed to cause a likely change in phenotype. (I. A. Adzhubei et al. 2010) We used the PolyPhen-2 analysis obtained by the Rhapsody web interface. Rhapsody enables automated pathogenicity evaluation that incorporates sequence coevolution data, as well as structural and dynamical features. (Ponzoni et al. 2020)

### 2.8. SIFT

SIFT (Sort Intolerant From Tolerant) is an algorithm predicting the impact of coding variants on protein. SIFT uses sequence homology to estimate the probability that an amino acid substitution will be deleterious to protein function. (Ng and Henikoff 2003) Starting with a given protein sequence, SIFT identifies evolutionarily related proteins and generates a corresponding multiple sequence alignment with the query sequence. At each alignment position, the algorithm evaluates the amino acid distribution and estimates the likelihood that a specific residue will be tolerated, assuming that the most common amino acid at that site is functionally acceptable. If this normalized probability falls below a predefined threshold, the variant is classified as potentially harmful or deleterious. (Sim et al. 2012) In contrast with PolyPhen-2 and AlphaMissense, which use the scale of pathogenicity from 0 (benign) to 1 (pathogenic), the SIFT score is a normalized probability of observing the new amino acid at that position and ranges from 0 to 1, where the value between 0 and 0.05 is predicted to affect protein function and can be described as deleterious. (Ng and Henikoff 2003) The SIFT algorithm uses a protein FASTA file as input for the analysis.

Python 3.9.13 using the os, json, pandas, seaborn, and matplotlib.pyplot libraries were used to process the results obtained from the AlphaMissense, SIFT, and PolyPhen-2 tools, as well as to create the figure for the article, unless otherwise stated. Three tools for predicting pathogenic MMuts were used to confirm our basic hypothesis of increased pathogenicity at sites functionally necessary for the protein. The three techniques described in Table 1 are presented with their features.

**Table 1:** Division of three MM pathogenicity prediction techniques by their features.

| Feature | AlphaMissense | SIFT | PolyPhen-2 |
| --- | --- | --- | --- |
| Principle | Utilizes deep learning approaches to predict variant effects based on sequence and structural information | Evaluates evolutionary conservation using multiple sequence alignments across species | Combines sequence- and structure-based features processed through a Naive Bayes classifier to predict functional impact |
| Data basis | Large-scale variant databases combined with structural models generated by AlphaFold | Cross-species sequence alignments reflecting evolutionary constraints | Human protein sequences, detailed structural annotations, and known variant datasets |
| Output | Provides a probability score of pathogenicity ranging from 0 to 1; higher values indicate a higher likelihood of pathogenicity | Generates a score between 0 and 1; variants with scores below 0.05 are predicted to be deleterious | Produces a qualitative classification (benign, possibly damaging, probably damaging, damaging) along with a confidence score from 0 to 1 |
| Strengths | Incorporates 3D structural context from AlphaFold predictions, enabling broad proteome coverage and strong generalization capability | A fast, simple, and interpretable method widely used in variant effect prediction | Effectively integrates sequence and structural data, widely adopted with proven performance, especially on known human variants |
| Limitations | Performance is dependent on the quality of AlphaFold structural models, which may be less reliable for rare or poorly characterized proteins | Does not include structural information, less accurate for proteins with low conservation or short length | Accuracy hinges on the training dataset, may misclassify rare or novel variants, and assumes certain structural relationships that might not always hold |

### 2.9. Evaluation based on experimental data

To evaluate the computationally predicted intolerance to substitution in GLUTs, we used data from the ClinVar database, a publicly accessible repository of more than three million human genetic variants. ClinVar classifies variants into three categories: germline classification, oncogenicity, and the clinical impact of somatic variants. (Landrum et al., 2025, 2016, 2014) For this study, we retrieved only missense variants in GLUT transporters classified as pathogenic or benign, with a minimum review status of one star. The full list of retrieved variants is shown in Table 02. We were able to obtain data of our preference only for the following proteins: GLUT1, GLUT2, GLUT5, GLUT6, GLUT7, GLUT8, GLU9, and GLU10.

These clinically observed classifications were then used as a benchmark to compare the predictions of three computational methods (AlphaMissense, PolyPhen-2, and SIFT), and the predictive accuracy of these methods was assessed using ROC-AUC analysis. GLUT3, GLUT4, GLUT11, GLUT12, GLUT13, and GLUT14 had no qualifying ClinVar variants and were therefore excluded.

## 3. Results

The main objective of this study was to verify the predicted pathogenicity of individual GLUT class protein components, assess the impact of MMuts on protein structure, and identify the components most affected by them. Firstly, we performed an analysis using Boltz-2 to get a first look at the protein structures. The protein structures were organized into the described groups. For the first and second groups, the structures were predicted to be outward-facing; for the third group, the predictions were predominantly occluded, with two—GLUT6 and GLUT13—inward-facing. The reasons for this deviation in the prediction could have a possible explanation. GLUT6 and GLUT13 show greater sequence divergence from the well-characterized Group I transporters used to benchmark structure-prediction models, potentially biasing Boltz-2 toward a less confidently sampled conformational state. Since no experimentally resolved structures currently exist for GLUT6 or GLUT13, we cannot presently determine whether this reflects a modeling limitation or a biologically meaningful difference in conformational preference.

The predicted structures were used to calculate the transport pathway—the pore—using MOLE and the binding site using PrankWeb, and to classify proteins as intracellular, extracellular, or transmembrane using DeepTMHMM. An overview of the structures is shown in Figure 2.

**Figure 2:**
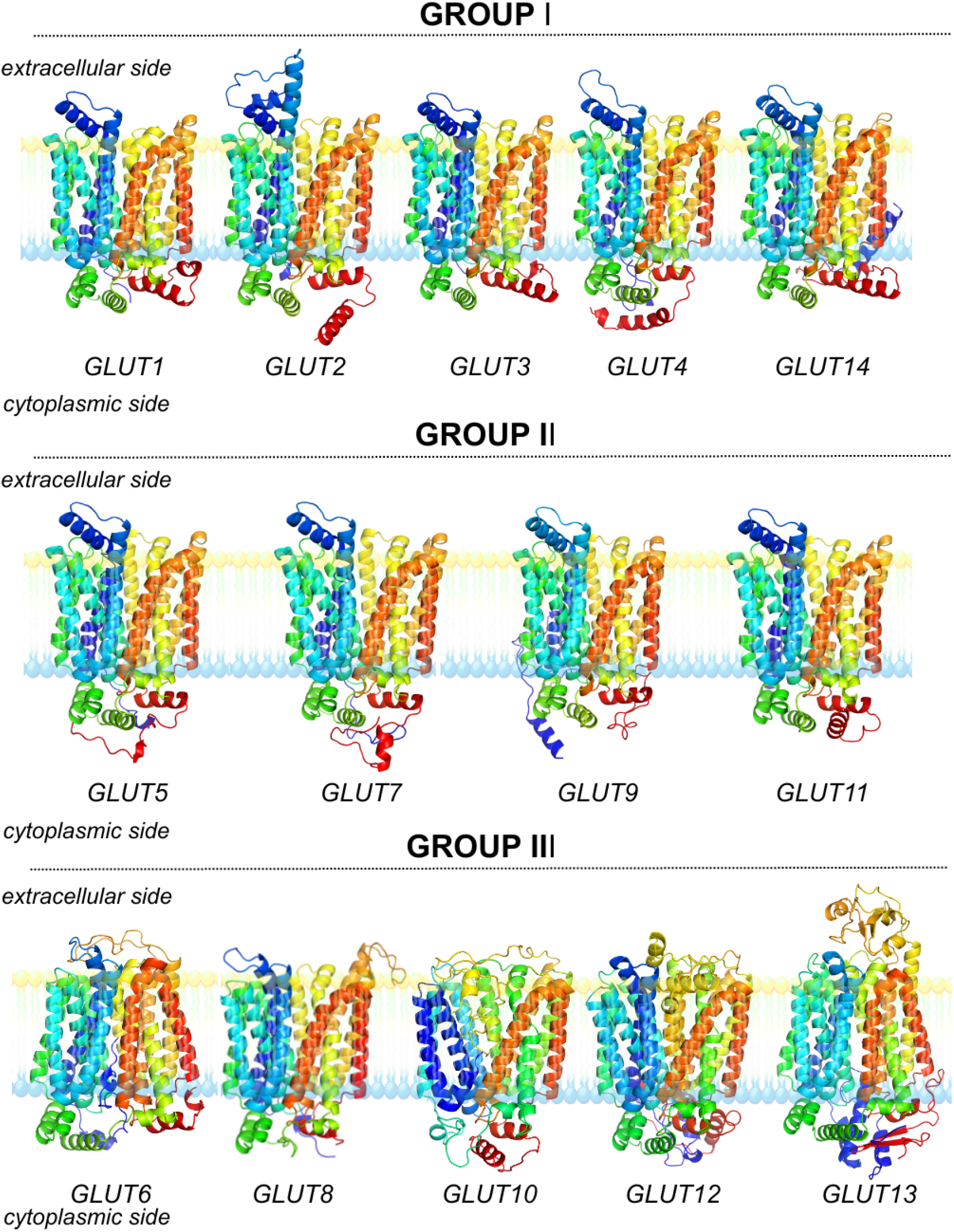
Predicted protein structures through Boltz-2. Protein models are embedded in membranes marked by a lipid bilayer separating the extracellular and cytoplasmic sides. Protein membrane coordination was assessed using DeepTMHMM. The coloring scale corresponds to sequence position (rainbow, N-to-C) in the palette, from blue to red.

### 3.1. The regions of higher intolerance to amino acid substitutions

We have analyzed the pathogenicity of 14 GLUT representatives using 3 prediction methods: AlphaMissense, PolyPhen-2 via the Rhapsody web interface, and SIFT via its web interface. The results for each prediction technique were recorded separately for each protein and then applied to its individual parts. We have separately analyzed the extracellular and intracellular parts, TM regions, binding sites, and lining residues along the transport pathway. Results (Figure 3) show a direct increase in pathogenicity towards the pore and binding site. It can be said that AlphaMissense, PolyPhen-2, and SIFT predicted similar trends for the regions. The computationally predicted intolerance to amino acid substitution is significantly higher here than in the transmembrane region overall.

**Figure 3:**
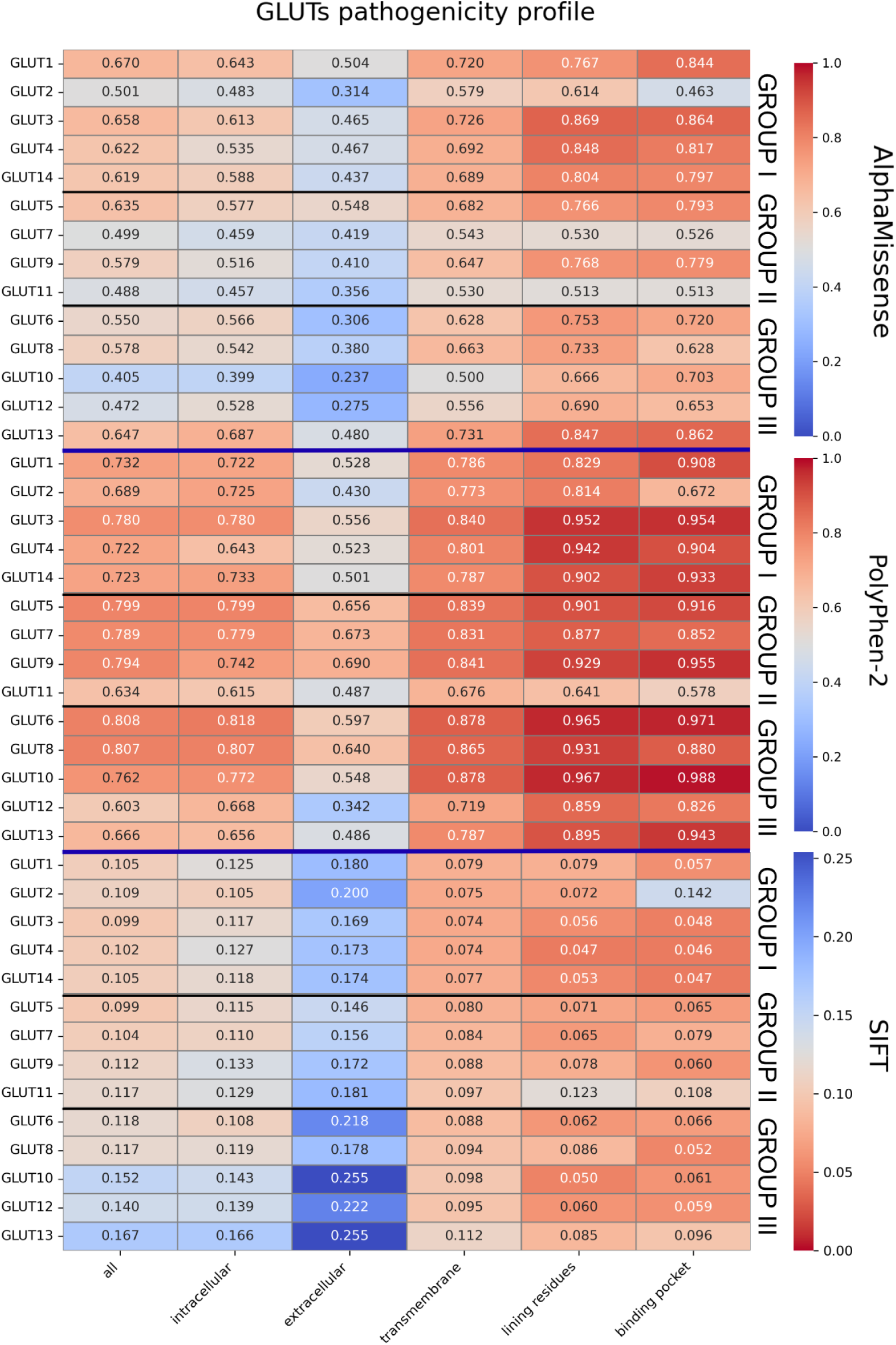
The average pathogenicity scores obtained from PolyPhen-2 (via Rhapsody), AlphaMissense, and SIFT are shown for seven different protein regions represented in the form of a heatmap. The AlphaMissense and PolyPhen-2 scales range from 0 (non-pathogenic/benign) to 1 (highly pathogenic/malignant). The SIFT method uses a different, reversed scale ranging from 1 (non-deleterious) to 0 (deleterious). Blue was used to mark benign mutations and red to mark highly pathogenic, malignant mutations.

In contrast, the TM regions were found to be more pathogenic than the extracellular and intracellular domains, consistent with Tordai et al. Finally, the extracellular domain was identified as the least affected by pathogenic MMuts. (Figure 4). Taken together, the convergence of three independent methods on the same regional hierarchy strengthens confidence that binding-site and pore-lining residues are genuinely more mutation-intolerant than residues in the extracellular domain of this transporter family.

**Figure 4:**
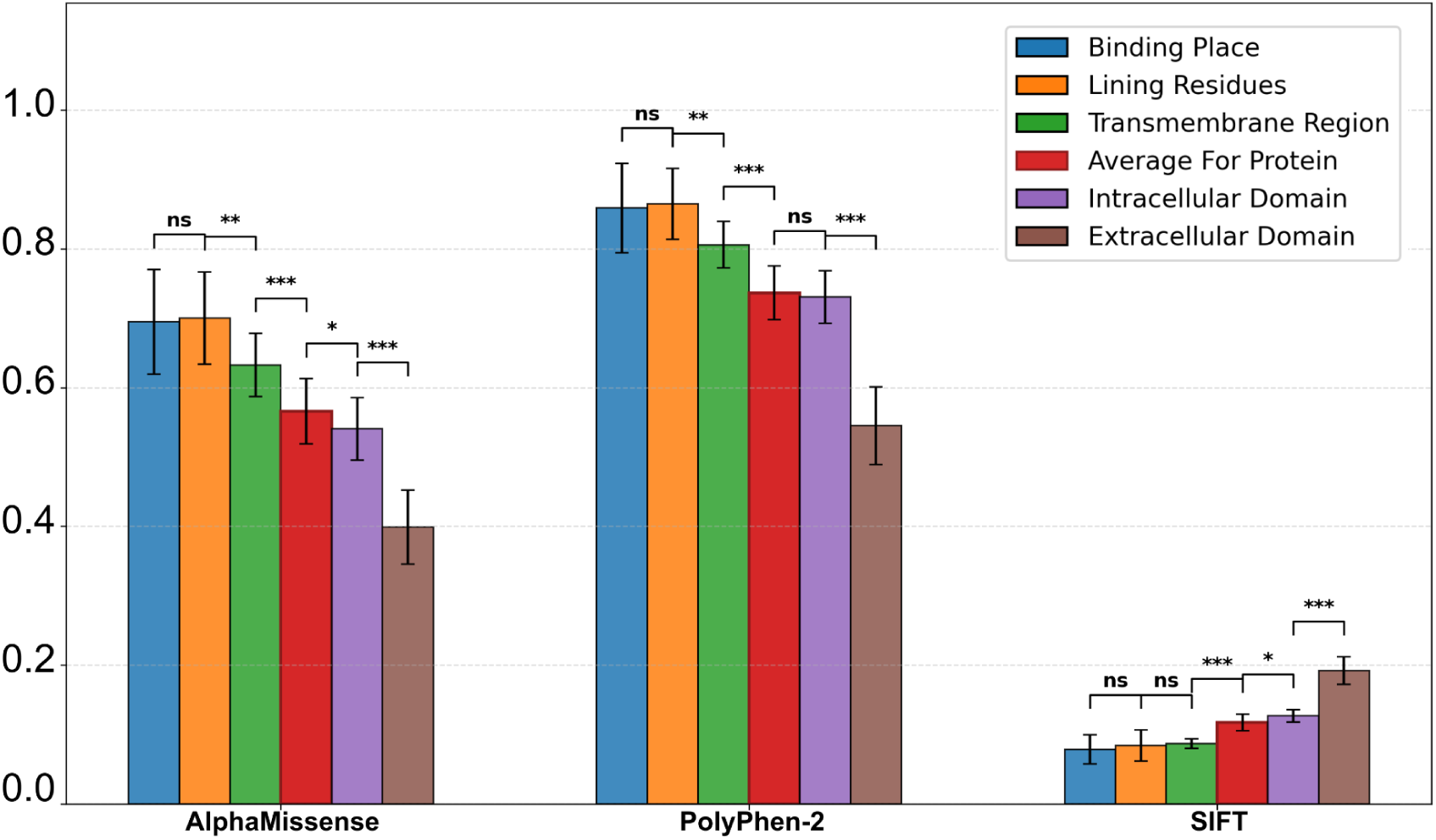
Statistical analysis of pathogenicity scores from PolyPhen-2, AlphaMissense, and SIFT for individual protein regions. The bars represent the mean scores with 95% confidence intervals. Asterisks above the bars denote statistical significance based on paired t-tests comparing pathogenicity as follows: Protein vs Transmembrane Part vs Intracellular Domain vs Extracellular Domain, and then Protein vs Lining Residues vs Binding Place (* p < 0.05, ** p < 0.005, *** p < 0.0005). All three methods consistently indicate an increase in pathogenicity from the general protein background towards the transmembrane part, with the most pronounced differences observed in AlphaMissense and PolyPhen-2. SIFT exhibits the same trend but uses an inverse scoring system, where lower values indicate higher pathogenicity. These findings emphasize that functionally critical regions, such as the pore and the binding site, are more sensitive to MMuts.

### 3.2. The transportational pathway pathogenicity

To verify the pathogenicity of the TM region, a structural alignment was generated using DALI and AlphaMissense data. Twelve areas marked as TM helices are clearly visible, along with the highest number of pathogenically predicted MMuts. These regions correspond to 12 helices passing through the membrane. Through this verification, we can confirm elevated predicted deleteriousness scores in the protein’s TM region, a pattern observed across all 14 proteins (Figure 5B). This also confirmed a high degree of pathogenicity in the helices surrounding the transport pathway and the binding cavity. This trend is consistent across all observed proteins. Moreover, a pathogenicity profile was created for each TM helix separately. The highly pathogenic TM helices can be identified as 1, 2, 4, 5, 7b, and 11, which directly border the internal transport pathway—the pore (Figure 5C). At the same time, however, some GLUTs exhibit an overall decrease in pathogenicity relative to others, even within highly pathogenic regions. These proteins are GLUT2, GLUT7, GLUT11, and GLUT10, which exhibited an overall low-pathogenicity profile in the initial analysis of individual regions. Conclusively, a pathogenicity profile based on AlphaMissense predictions shows a similar pattern across all GLUT family members. It is also evident that GLUT1 is a representative of highly pathogenic proteins, followed by GLUT2 with a lower pathogenicity profile, and GLUT10 as one of the GLUTs with the lowest pathogenicity profile (Figure 5A). Pathogenic non-synonymous amino acid substitutions are similarly distributed across all protein regions, showing the same trend: increased pathogenicity in the vicinity of the central cavity (binding site) and the pore. However, an overall decrease in the pathogenicity profile within the protein structure is specific to the individual.

**Figure 5:**
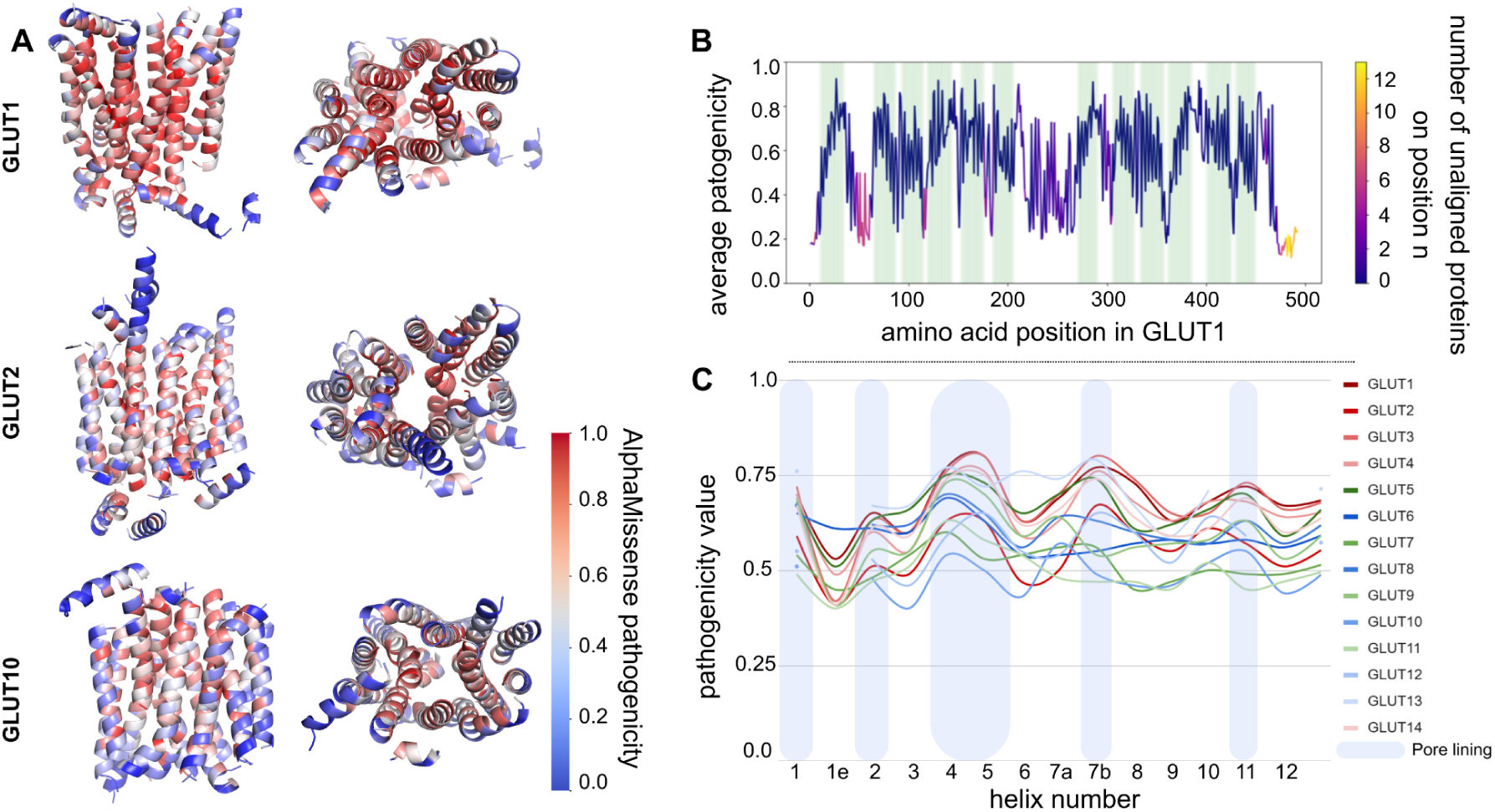
Pathogenicity analysis of individual transmembrane helices. **A**: Pathogenicity profile of non-synonymous substitutions (AlphaMissense) in three representative GLUT proteins, shown in side (left) and top (right) views. GLUT1 shows high pathogenicity throughout, while GLUT10 and GLUT2 show progressively lower pathogenicity, with benign substitutions predominating. In all three proteins, pathogenicity increases radially toward the central transport channel. Color scale: blue (benign) to red (pathogenic). **B**: Overlap of highly pathogenic sites shown on a DALI-based structural alignment of 13 GLUT family proteins relative to GLUT1, with transmembrane helices marked by green stripes. **C**: Pathogenicity profile of transmembrane helices across GLUT proteins. Helices 1, 2, 4, 5, 7b, and 11 show the highest pathogenicity and line the pore and binding cavity; helices 1e, 3, 6, 8, 9, and 10 show the lowest pathogenicity and form the outer, non-transport-facing rim of the transmembrane region.

### 3.3. Comparison to experimental data from ClinVar

The ROC-AUC analysis (Figure 6B) provides direct validation of the predictor-based findings described above. AlphaMissense achieved the highest discriminative performance (AUC = 0.91), followed by SIFT (AUC = 0.83) and PolyPhen-2 (AUC = 0.83), confirming that the pathogenicity trends observed using AlphaMissense throughout this study are the most reliable of the three methods tested, though all three retain reasonable predictive accuracy against clinically observed variants. The helix-level ClinVar analysis (Figure 6C) largely supports the predictor-based helix pathogenicity profile described in Figure 5C: helices 4, 5, 9, and 11 show a marked excess of pathogenic over benign variants, consistent with their identification as high-pathogenicity, pore-lining helices in the AlphaMissense-based analysis. Helix 1, however, shows a more balanced ratio in the ClinVar data despite being predicted as highly pathogenic. This discrepancy is likely attributable to the limited size of the ClinVar dataset: only 140 variants across all 14 GLUT proteins met our classification and review-status criteria (Table 2), and this constraint is more pronounced at the level of individual helices, where the number of qualifying variants per helix is correspondingly small.

**Figure 6:**
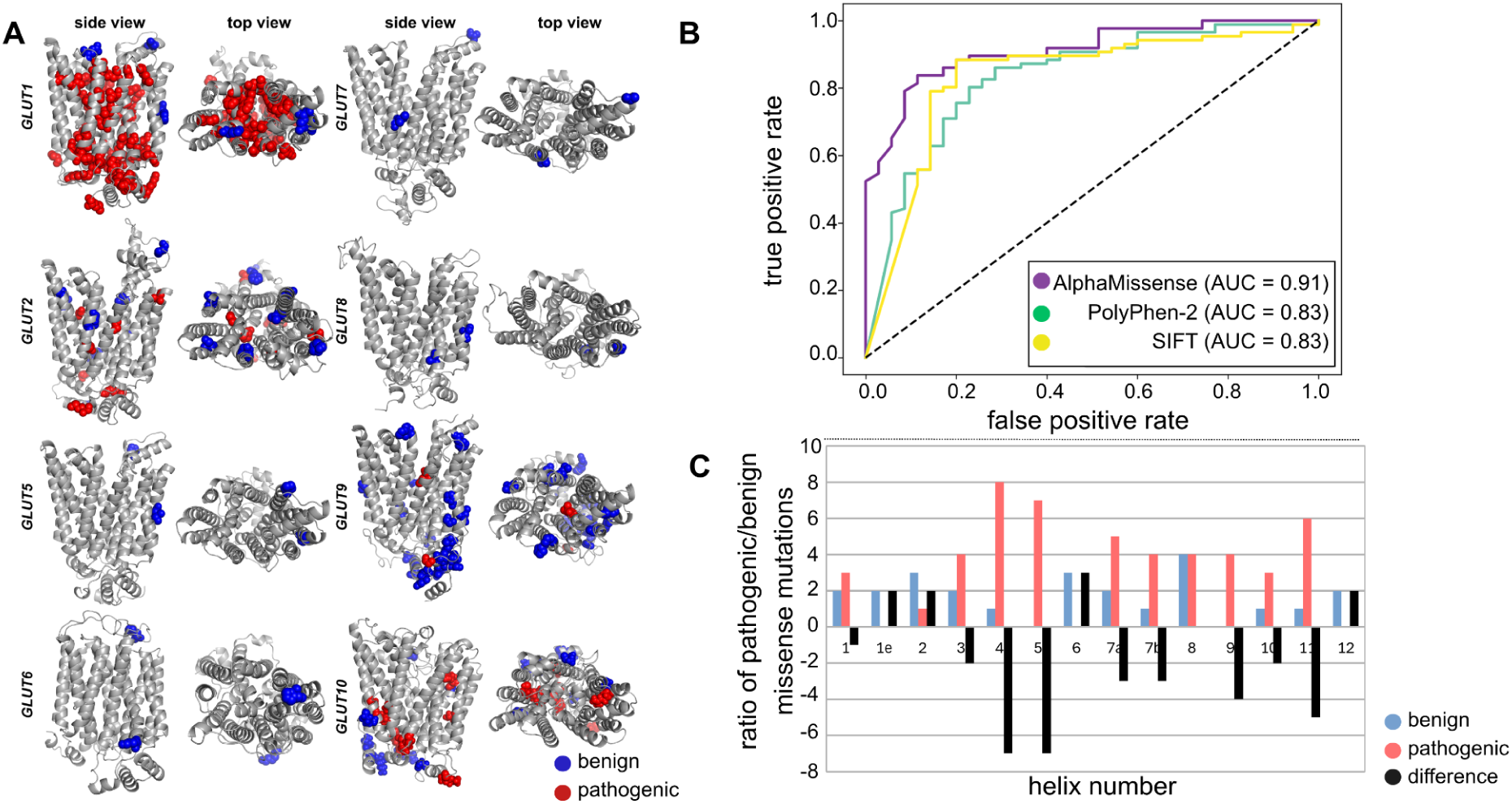
Experimental validation of predicted pathogenicity using ClinVar variant data. **A:** Structural mapping of ClinVar-classified benign (blue) and pathogenic (red) missense variants onto the modeled structures of eight GLUT proteins (GLUT1, 2, 5, 6, 7, 8, 9, 10), shown in side and top views. GLUT1 and GLUT10 show a marked concentration of pathogenic variants, whereas GLUT5, GLUT6, GLUT7, and GLUT8 show predominantly benign variants, consistent with the low number of ClinVar entries available for these proteins (see Table 2). **B:** ROC curves comparing the performance of three pathogenicity prediction methods — AlphaMissense, PolyPhen-2, and SIFT — against pathogenic and benign missense variants retrieved from ClinVar. AlphaMissense showed the highest discriminative performance (AUC = 0.91), followed by SIFT (AUC = 0.83) and PolyPhen-2 (via Rhapsody) (AUC = 0.83). **C:** Ratio of pathogenic to benign ClinVar-classified missense variants across individual transmembrane helices. Helices 4, 5, 9, and 11 show a strong excess of pathogenic variants, while helices 1, 6, and 12 show a more balanced or benign-skewed distribution, indicating uneven tolerance to substitution across the transmembrane region.

**Table 2:** Number of missense variants retrieved from ClinVar for eight human GLUT transporters, classified as pathogenic/likely pathogenic or benign/likely benign, with a minimum review status of one star. Only GLUT proteins with at least one qualifying variant are shown. Totals for each classification and protein are indicated.

|  | **Pathogenic/**<br>**Likely pathogenic** | **Benign/**<br>**Likely benign** | **Total** |
| --- | --- | --- | --- |
| *GLUT1* | 71 | 3 | 74 |
| *GLUT2* | 7 | 9 | 16 |
| *GLUT5* | 0 | 2 | 2 |
| *GLUT6* | 0 | 2 | 2 |
| *GLUT7* | 0 | 2 | 2 |
| *GLUT8* | 0 | 2 | 2 |
| *GLUT9* | 2 | 23 | 25 |
| *GLUT10* | 9 | 8 | 17 |
| **Together** | **89** | **51** | **140** |

The structural mapping of ClinVar variants (Figure 6A) is consistent with the AlphaMissense-based structural profile (Figure 5A) for most proteins; GLUT1 shows a dense concentration of pathogenic amino acid changes throughout its structure, consistent with its identification as the most pathogenic GLUT family member in the predictor-based analysis. However, GLUT10 shows a notable cluster of pathogenic ClinVar variants concentrated around its transport pathway, which contrasts with its classification as one of the lowest-pathogenicity proteins in the AlphaMissense-based profile (Figure 5A). As with helix 1, this discrepancy likely reflects the restricted number of ClinVar entries available per protein (Table 2) rather than a true divergence between predicted and observed pathogenicity. The overall sparseness of the ClinVar dataset should therefore be considered when interpreting protein- and helix-level agreement between predicted and clinically observed pathogenicity, even though these values represent the full set of variants meeting our classification criteria at the time of analysis.

## 4. Discussion

The human GLUT family, comprising 14 facilitative transporters (*SLC2A*), acts as the primary gateway for glucose and other hexoses, ensuring that the diverse energy demands of tissues are met through regulated facilitated diffusion. (Navale and Paranjape 2016)

In the major facilitator superfamily (MFS) fold typical of GLUTs, the N-terminal (TM1-6) and C-terminal (TM7-12) bundles form a central cavity. Crucial interactions for substrate recognition, such as hydrogen bonds with residues in TM1 and TM7, occur deep within this transmembrane region. (Khandelwal et al., 2025) Pathogenic MMuts cluster mostly in this region, toward the center of the protein structure, which houses the substrate-binding pocket and the translocation pathway. TM7 often acts as a “lid” or part of a selectivity filter for the translocation pore. The transporting motion requires these central transmembrane bundles to reorient precisely to move the substrate. (Cheeseman, 2008; Lee et al., 2024) Studies on GLUT1 and GLUT9 (*SLC2A9*) demonstrate that mutations in central segments (such as TM7) directly alter kinetic parameters. (Khandelwal et al., 2025; Wong et al., 2007) TM7, together with TM1, TM2, TM4, TM5, and TM11, which form the inner lining of the transport pathway, showed high pathogenicity. Non-synonymous amino acid substitutions in the central core would likely disrupt the high-affinity binding sites or the conformational flexibility needed to switch between inward- and outward-open states.

The pathogenicity trends described above are further supported by comparison with clinically observed variants from ClinVar. AlphaMissense showed the strongest agreement with clinical classifications (AUC = 0.91), outperforming PolyPhen-2 and SIFT, and the helix- and structure-level distribution of pathogenic and benign ClinVar variants broadly recapitulated the predicted pathogenicity patterns, most notably the concentration of pathogenic substitutions around the pore-lining helices (TM1, TM2, TM4, TM5, TM7, TM11). The elevated pathogenicity of GLUT1 compared with other family members is also evident. This concordance between predicted and clinically observed pathogenicity supports the biological plausibility of the structural interpretations discussed below, though the relatively small number of ClinVar entries meeting our inclusion criteria (140 variants across 14 proteins) limits the resolution of this validation at the level of individual helices and lower-abundance GLUT isoforms.

The observed systematically lower values in pathogenic mutations in Groups I to III can be discussed in the context of physiological essentiality and missense tolerance.

Group I transporters (GLUT1-4,14) are the best-characterized and most physiologically critical isoforms. (Navale and Paranjape 2016; Chadt and Al-Hasani 2020) Proteins in this group show high pathogenicity values, except for GLUT2, which stands out with a low pathogenicity profile. GLUT2 is a bidirectional transporter essential for glucose sensing in the liver, pancreas, and kidneys. (Holman, 2020) Mutations here often lead to severe pathologies like Fanconi-Bickel syndrome, which is an autosomal recessive pathology. (Grünert et al., 2012) Recessive genes typically show a lower pathogenicity profile because the protein is often functionally tolerant of variation in one allele. (Huang, 2020) GLUT2 is also described as a low-affinity, high-capacity isoform, engineered to work at high glucose concentrations. (Carreño et al., 2021) Then the structural precision of its binding pocket may be less sensitive to minor missense mutations than that of transporters that must capture glucose at very low concentrations, such as GLUT1. (Bourque et al., 2021) Lastly, evolutionary conservation may play a role. While GLUT1 is highly conserved across species to maintain basal respiration (Bourque et al., 2021), GLUT2 has undergone greater evolutionary diversification due to its varied roles in the liver, kidney, and intestine, and thus probably shows a lower pathogenicity profile.

Group II transporters often exhibit more localized expression or overlap in function with other hexose transporters. (Cheeseman, 2008; Schürmann, 2008) GLUT7 and GLUT11 facilitate the transport of both glucose and fructose. The notable difference in the mutation profile may reflect that their physiological roles are partially compensated for by other Class II members, such as GLUT5. (Song et al., 2023) Notably, GLUT9 shows a ClinVar-derived variant profile skewed strongly toward benign classifications (2 pathogenic/likely pathogenic vs. 23 benign/likely benign; Table 2), despite AlphaMissense predicting relatively high pathogenicity for several of its pore-lining helices (Figure 5C). GLUT9 is functionally distinct within Group II as a urate transporter with a comparatively narrow, tissue-restricted physiological role rather than a broad role in glucose homeostasis (Doblado and Moley, 2009), which may make it more tolerant of substitutions outside residues directly required for urate recognition and transport. This pattern parallels the GLUT2 case in Group I, where a specialized, low-affinity transport function is associated with reduced predicted pathogenicity, and suggests that functional specialization, rather than family-wide essentiality alone, may be an important factor shaping missense tolerance across GLUT groups. As with GLUT10, however, the relatively small number of ClinVar entries available for GLUT9 warrants some caution in drawing strong conclusions from this comparison.

Group III transporters show the lowest pathogenicity profile for GLUT10. GLUT10 is structurally distinct from other GLUTs due to the absence of the PESPR domain. (Németh et al., 2016) While GLUT10 deficiency causes Arterial Tortuosity Syndrome (ATS), its role in systemic glucose homeostasis is less ubiquitous than that of Class I transporters. (Esmel-Vilomara et al., 2023; Németh et al., 2016) Interestingly, this low overall predicted pathogenicity contrasts with the ClinVar-derived structural mapping, which shows a concentration of pathogenic variants around the GLUT10 transport pathway despite its low genome-wide predicted pathogenicity. This may indicate that while GLUT10 tolerates substitution across most of its structure, the residues directly involved in transport retain strong functional constraint — consistent with a physiological role that is narrower but still critical where it occurs, such as in the integrity of vascular connective tissue. Alternatively, given the modest number of ClinVar entries available for GLUT10 (Table 2), this pattern may partly reflect ascertainment bias toward reporting variants associated with the known clinical phenotype (ATS) rather than a complete, unbiased sampling of tolerated versus intolerant residues.

## 5. Limitations

The limitations of our study should be considered when interpreting these findings. First, structural models for 9 of the 14 GLUT proteins were generated computationally using Boltz-2 in the absence of experimentally resolved structures. While Boltz-2 provides a reasonable basis for family-wide comparison, model inaccuracies cannot be ruled out, particularly for Group III transporters, for which structural and functional characterization remains limited. Second, the ClinVar-based validation was constrained by the relatively small number of qualifying variants; six GLUT family members had no qualifying entries and therefore could not be included in this validation step. The uneven distribution of pathogenic and benign variants across the remaining eight proteins further limits the resolution of protein- and helix-level comparisons. This may reflect ascertainment bias in clinical variant reporting rather than a complete, unbiased sampling of tolerated and intolerant residues. Finally, this study relies entirely on computational prediction and clinically reported variant classifications; it does not include direct functional or experimental validation (e.g., transport assays, structural determination of variant proteins), and the proposed biological explanations for group- and protein-level differences in pathogenicity should therefore be regarded as plausible interpretations consistent with the available evidence rather than directly demonstrated mechanisms.

## 6. Conclusion

We conclude that missense mutations are not randomly distributed across GLUT family proteins but instead cluster within the substrate-binding cavity and the translocation pathway. This pattern was independently supported by comparison with clinically observed variants from ClinVar: AlphaMissense showed the strongest concordance with clinical classifications (AUC = 0.91) among the three prediction methods tested, and the structural distribution of clinically reported pathogenic variants largely mirrored the predicted pathogenicity landscape. It seems that the transport pathway is the primary hub for pathogenic missense mutations, a conclusion now grounded in both computational predictions and clinical variant data rather than in predictions alone. On the other hand, this study observed differences in pathogenicity among some GLUTs across all three groups, and in some cases, such as GLUT10, the clinical data suggested that functional constraint may be more localized to the transport pathway than the overall predicted pathogenicity profile implied. This highlights that predicted and observed pathogenicity, while broadly concordant, are not always identical and should be interpreted together rather than separately. In conclusion, the evolutionary landscape of glucose transport is not uniform but shaped by varying degrees of physiological essentiality and functional redundancy across the three GLUT groups, a picture supported by convergent evidence from structural, evolutionary, and clinical variant data.

## Data Availability Statement

The input data for the model and the predictions used in this study are available at https://github.com/DominikMartinat/GLUT_pathogenicity_analysis. The same workflow and resulting datasets are also available on Zenodo https://doi.org/10.5281/zenodo.22011238.

## Author contributions

Nina Kadášová (Formal analysis, Methodology, Visualization, Writing—original draft), Anna Špačková (Methodology, Visualization, Writing—original draft), Dominik Martinát (Methodology, Visualization, Writing—original draft), Ivana Hutařová Vařeková (Methodology, Visualization), and Karel Berka (Methodology, Project administration, Supervision, Writing—original draft).

## Funding and additional information

This work has been supported by the Palacký University Olomouc [IGA_PrF_2026_002 to N.K., A.Š., and D.M.], and ELIXIR CZ research infrastructure projects (MEYS) [LM2023055 - ELIXIR CZ and 8K0208 - ELIXIR IMPACT] to N.K., A.Š., D.M., I.H.V., and K.B.

